# Genome-wide association study of social relationship satisfaction: significant loci and correlations with psychiatric conditions

**DOI:** 10.1101/196071

**Authors:** Varun Warrier, Thomas Bourgeron, Simon Baron-Cohen

**Affiliations:** Autism Research Centre, Department of Psychiatry, University of Cambridge, Cambridgeshire, United Kingdom; 23andMe Inc., Mountain View, CA; Institut Pasteur, Human Genetics and Cognitive Functions Unit, Paris, France; CNRS UMR 3571: Genes, Synapses and Cognition, Institut Pasteur, Paris, France; Université Paris Diderot, Sorbonne Paris Cité, Human Genetics and Cognitive Functions, Paris, France; CLASS Clinic, Cambridgeshire and Peterborough NHS Foundation Trust (CPFT), Cambridgeshire, United Kingdom

## Abstract

Dissatisfaction in social relationships is reported widely across many psychiatric conditions. We investigated the genetic architecture of family relationship satisfaction and friendship satisfaction in the UK Biobank. We leveraged the high genetic correlation between the two phenotypes (r_g_ = 0.87±0.03; P < 2.2x10^-16^) to conduct multi-trait analysis of Genome Wide Association Study (GWAS) (N_effective_ family = 164,112; N_effective_ friendship = 158,116). We identified two genome-wide significant associations for both the phenotypes: rs1483617 on chromosome 3 and rs2189373 on chromosome 6, a region previously implicated in schizophrenia. eQTL and chromosome conformation capture in neural tissues prioritizes several genes including *NLGN1*. Gene-based association studies identified several significant genes, with highest expression in brain tissues. Genetic correlation analysis identified significant negative correlations for multiple psychiatric conditions including highly significant negative correlation with cross-psychiatric disorder GWAS, underscoring the central role of social relationship dissatisfaction in psychiatric diagnosis. The two phenotypes were enriched for genes that are loss of function intolerant. Both phenotypes had modest, significant additive SNP heritability of approximately 6%. Our results underscore the central role of social relationship satisfaction in mental health and identify genes and tissues associated with it.

## Introduction

Social relationship satisfaction is one contributor to subjective wellbeing, alongside other domains such as occupational, financial, and health satisfaction^6^. Difficulties in forming and maintaining social relationships are reported widely in psychiatric conditions^1–4^. Positive social relationship satisfaction can both reduce the risk and ameliorate the severity of a psychiatric condition. In the DSM-5^5^, difficulties in social functioning is one of the criteria for diagnosing conditions such as autism, anorexia nervosa, schizophrenia, and bipolar disorder. However, little is known about the genetic architecture of social relationship satisfaction, and if social relationship dissatisfaction genetically contributes to risk for psychiatric conditions. To our knowledge, there is no twin study that has focussed on twin heritability of social relationship satisfaction. A few studies have investigated the heritability of subjective wellbeing and have identified heritability estimates between 38 – 50%^7,8^.

Here we report on two genome-wide association studies (GWAS) of social relationship satisfaction in the UK Biobank. We focus on friendship satisfaction and family relationship satisfaction, measured using two questionnaires. We leverage the high genetic correlation between the two phenotypes to: 1. Identify the genetic architecture of friendship and family relationship satisfaction; 2. Identify the genetic correlation between these two phenotypes and specific psychiatric conditions and psychological phenotypes; 3. Prioritize associated genes, and enriched gene sets and tissues for the two phenotypes; 4. Quantify cross-phenotype polygenic predictive power of the two phenotypes.

We identified two genetic loci associated with the phenotypes. We further identified significant genetic correlations with several psychiatric conditions, and demonstrated an enrichment for brain tissues, and conclude by prioritizing genes and gene sets for further investigation.

## Methods

### Phenotypes and participants

Participants were individuals from the UK Biobank. Participants were asked “In general, how satisfied are you with your family relationships?” and “In general, how satisfied are you with your friendships?” Participants could choose one of eight options: “Extremely happy”, “Very happy”, “Moderately happy”, “Moderately unhappy”, “Very unhappy”, “Extremely unhappy”, “Do not know”, and “Prefer not to answer”. We excluded individuals who responded “Do not know”, or “Prefer not to answer” The phenotypes were coded in three waves. We combined responses from three waves, and removed participants who had responded in more than one wave (N = 178,675 for family and N = 178,721 for friendship). Responses were recoded from 1 to 6 with 1 being “Extremely unhappy”, and 6 being “Extremely happy”. We removed participants who were not genotyped, who were not of “British ancestry”, who were outliers for heterozygosity, and whose reported sex did not match their genetic sex. This left us with N = 139,603 for family, and N = 139,826 for friendship. Finally, we removed related individuals from this list resulting in a total of N = 134,681 (family) and N = 134,941 (friendship) unrelated individuals for further analyses. A total of 131,790 individuals were included in both the analyses. At the time of recruitment, participants were between 40 – 69 years of age. All participants provided informed consent to participate in the UK Biobank. In addition, we obtained ethical approval from the University of Cambridge Human Biology Research Ethics Committee (HBREC) to use de-identified data from the UK Biobank and ALSPAC for this study. ALSPAC is Avon Longitudinal Study of Parents and Children^27^ which was used in a polygenic risk score analysis (see below).

### Genetic analyses

Details of the UK Biobank genotyping, imputation, and quality control procedures is available elsewhere^9^. Genetic association was conducted using Plink 2.0^10^ (https://www.cog-genomics.org/plink/2.0/). We included sex, year of birth, genotyping batch and the first 40 genetic principal components as covariates. We excluded SNPs that failed Hardy-Weinberg Equilibrium (P < 1x10^-6^), had a minor allele frequency < 0.01, and had a per-SNP genotyping rate < 95%, and imputation INFO < 0.1. We further excluded SNPs not in the Haplotype Reference Consortium (http://www.haplotype-reference-consortium.org/)^11^. We excluded individuals who were ancestry outliers, and had per-individual genotyping rate < 90%.

To increase the effective sample size, we leveraged the high genetic correlation between the two phenotypes to conduct MTAG^12^ (or multi-trait analysis of GWAS) (https://github.com/omeed-maghzian/mtag/). MTAG is an extension of the standard inverse variance weighted meta-analysis that considers the genetic correlation between phenotypes to estimate phenotype-specific effects. Our phenotypes are excellent for MTAG given both the high genetic correlation between the two phenotypes and the similar mean chi-square. We do not expect the estimates to be biased.

*Clumping* of the independent SNPs was conducted using Plink^10^, using an r^2^ of 0.6. *Winner’s curse correction* was conducted using FIQT, which is an FDR based inverse quantile transformation^13^.

*Genetic correlations* were conducted using LD score regression (LDSR)^14,15^ (https://github.com/bulik/ldsc/wiki). We used north-west European population LD scores, did not constrain the intercept and used the pre-MTAG summary statistics to obtain unbiased genetic correlations. We ran correlations with psychiatric and psychological phenotypes **(Supplementary Table 1)**. Summary scores were obtained from the PGC (http://www.med.unc.edu/pgc/results-and-downloads), the SSGAC (https://www.thessgac.org/), and the CTG (https://ctg.cncr.nl/software/summary_statistics). In addition, summary statistics for systemizing, self-reported empathy^16^, and cognitive empathy^17^ were obtained from 23andMe, Inc. Summary statistics for the iPsych autism replication dataset were obtained from the iPsych autism team. Given the correlation between the phenotypes, we identified significant genetic correlations using a Benjamini-Hochberg FDR < 0.05, taking into account all the tests conducted for both the phenotypes combined.

*Heritability analyses* were conducted using both LDSR (all participants)^14^ and GCTA-GREML^18^ (for computational efficiency, this was conducted on one-fifth of the total participants, N = 26,000). To keep the methods as identical as possible, GCTA heritability was calculated after including sex, age, batch and the first 40 genetic principal components as covariates. Bivariate genetic correlations between the two phenotypes were conducted using both LDSR and GCTA.

*Functional annotation* was performed on the MTAG-GWAS datasets. We used FUMA^19^ (http://fuma.ctglab.nl/) to identify eQTLs and chromatin interactions for the genome-wide significant results. eQTLS were identified using data from BRAINEAC and GTEx^20^ brain tissues. Chromatin interactions were identified using Hi-C data from the dorsolateral prefrontal cortex, hippocampus, and neural progenitor cells. Significant mapping was identified at a Benjamini-Hochberg FDR < 0.05. Gene-based association analyses was conducted using MAGMA^21^ within FUMA, and significant genes were identified using a Bonferroni-corrected threshold < 0.05 for each phenotype, as provided within FUMA.

We conducted partitioned heritability^22^ analyses using the MTAG-GWAS dataset using the baseline categories and additional categories: genes that are loss-of-function intolerant^23^, and genes with brain specific annotation. Significant enrichments were identified using a Benjamini-Hochberg FDR < 0.05 applied to both the phenotypes combined.

To identify phenotype relevant tissues based on gene expression, we conducted partitioned heritability analyses applied to tissue-specific genes^24^. We focussed on GTEx consortium based gene expression from 13 brain regions (Cortex, Anterior cingulate cortex, frontal cortex, cerebellum, cerebellar hemisphere, putamen, nucleus accumbens, caudate, substantia nigra, hippocampus, spinal cord, amygdala, and hypothalamus). Results were significant at a Benjamini-Hochberg FDR < 0.05. In addition, gene expression analyses for general tissue types and specific tissue types were also conducted using FUMA. These two methods are not identical. In partitioned heritability, the top 10% of gene with tissue specific expression (focal brain region vs non-focal brain region) were included with a 100kb window on either side of the transcribed region. In FUMA, all genes are tested using a linear regression framework. Here, the expression of genes in a particular tissue is regressed against the gene Z-values with the average expression of the gene in all tissues included alongside usual covariates (gene length, SNP density etc). Thus, while FUMA tests all the genes, partitioned heritability tests only the top decile.

Cell specific enrichment^25^ was conducted for three cell types (Neurons, astrocytes and oligodendrocytes) using both LDSR partitioned heritability and MAGMA gene set enrichment. Results were FDR corrected (P < 0.05). For the gene-set enrichment with MAGMA, we used the top 500 cell-type specific genes for the three cell types. For oligodendrocytes, we used clustered both newly formed and myelinating oligodendrocytes into one group. The cell-specific genes can be downloaded here: https://web.stanford.edu/group/barres_lab/brain_rnaseq.html.

*Polygenic regression analyses* using the MTAG-GWAS results were conducted using PRSice^26^ to generate polygenic scores. Regression analysis was conducted in R version 3.3.3. Polygenic scoring was conducted in approximately 5,600 phenotyped and unrelated individuals from the Avon Longitudinal Study of Parents and Children^27^. We considered total scores from three parent-reported child-based questionnaires, collected between the ages of 7 and 10: the Children’s Communication Checklist (CCC)^28^, the Social and Communication Disorder Checklist^29^, and the Strengths and Difficulties Questionnaire (SDQ)^30^. These three questionnaires are widely used to assess difficulties in child development. In addition, we also tested five subdomains of the SDQ (prosocial behaviour, hyperactivity, emotional difficulties, peer difficulties and conduct problems). Prosocial behaviour and CCC were reverse coded to indicate difficulties in prosocial behaviour and communication. Polygenic score regression was conducted using a negative binomial model (MASS package in R: https://cran.r-project.org/web/packages/MASS/MASS.pdf) as this provided the best fit. Sex and the first two genetic principal components were included as covariates. Incremental variance explained due to polygenic scores was calculated using Nagelkerke’s pseudo-R^2^. Further details of the ALSPAC cohort are provided in the **Supplementary Note**.

## Results

### Phenotypic distributions

Both phenotypes had a unimodal, near-normal distribution (**Supplementary Figure 1**). The Spearman’s rank correlation between friendship and family relationship satisfaction was only 0.52 (P < 2.2x10^-16^) (**Supplementary Figure 2**), suggesting that they are not identical phenotypes. This allowed us to investigate the genetic architecture of the two phenotypes separately. Both sex and age were significantly associated with the two phenotypes, though the effect was small (**Supplementary Figures 3 and 4**).

**Figure 1:**
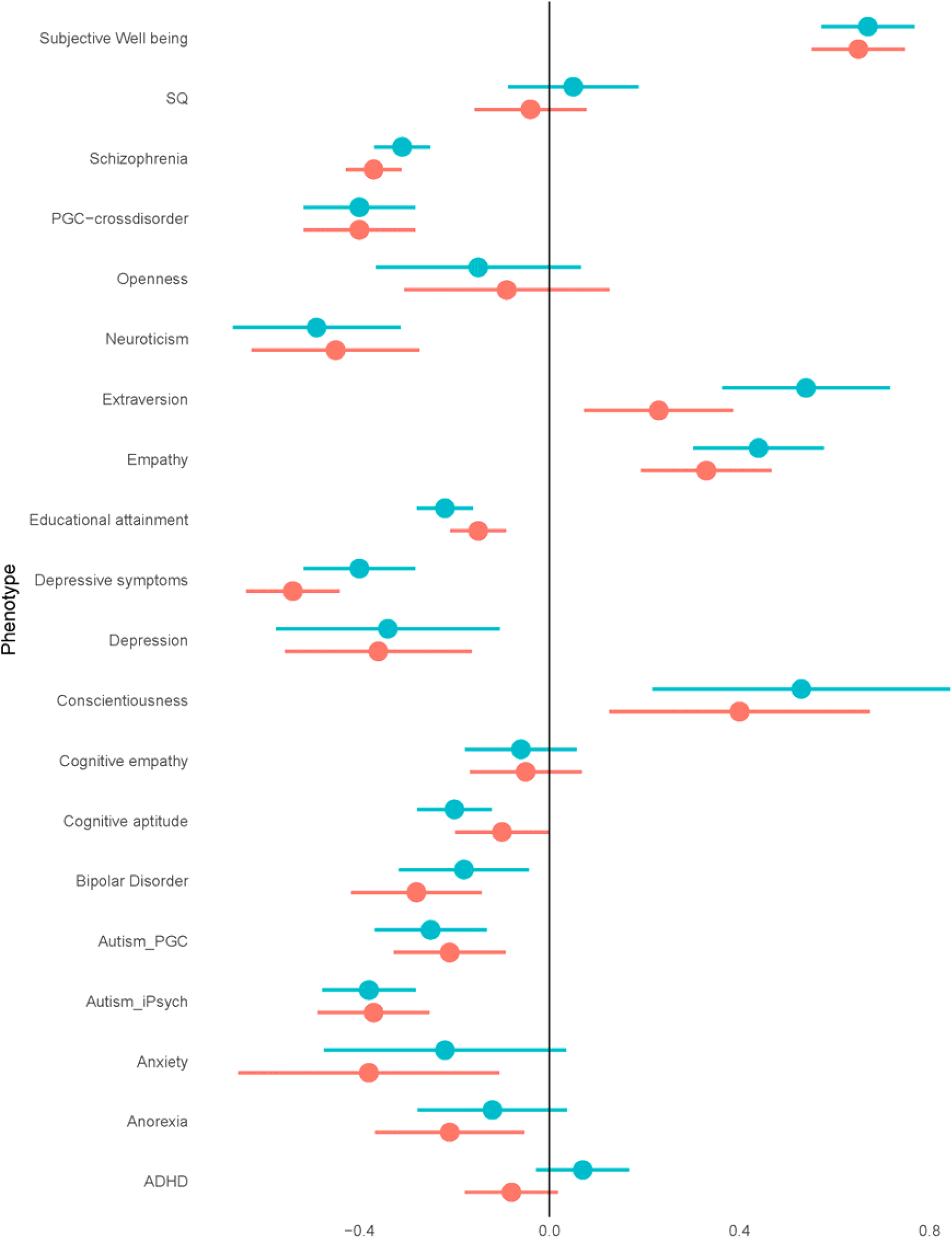
Genetic correlations. Genetic correlations and 95% confidence intervals provided for the two phenotypes (pink is family relationship satisfaction, blue is friendship satisfaction). Phenotypes tested are on the y axis. Genetic correlation is on the x axis.

**Figure 2:**
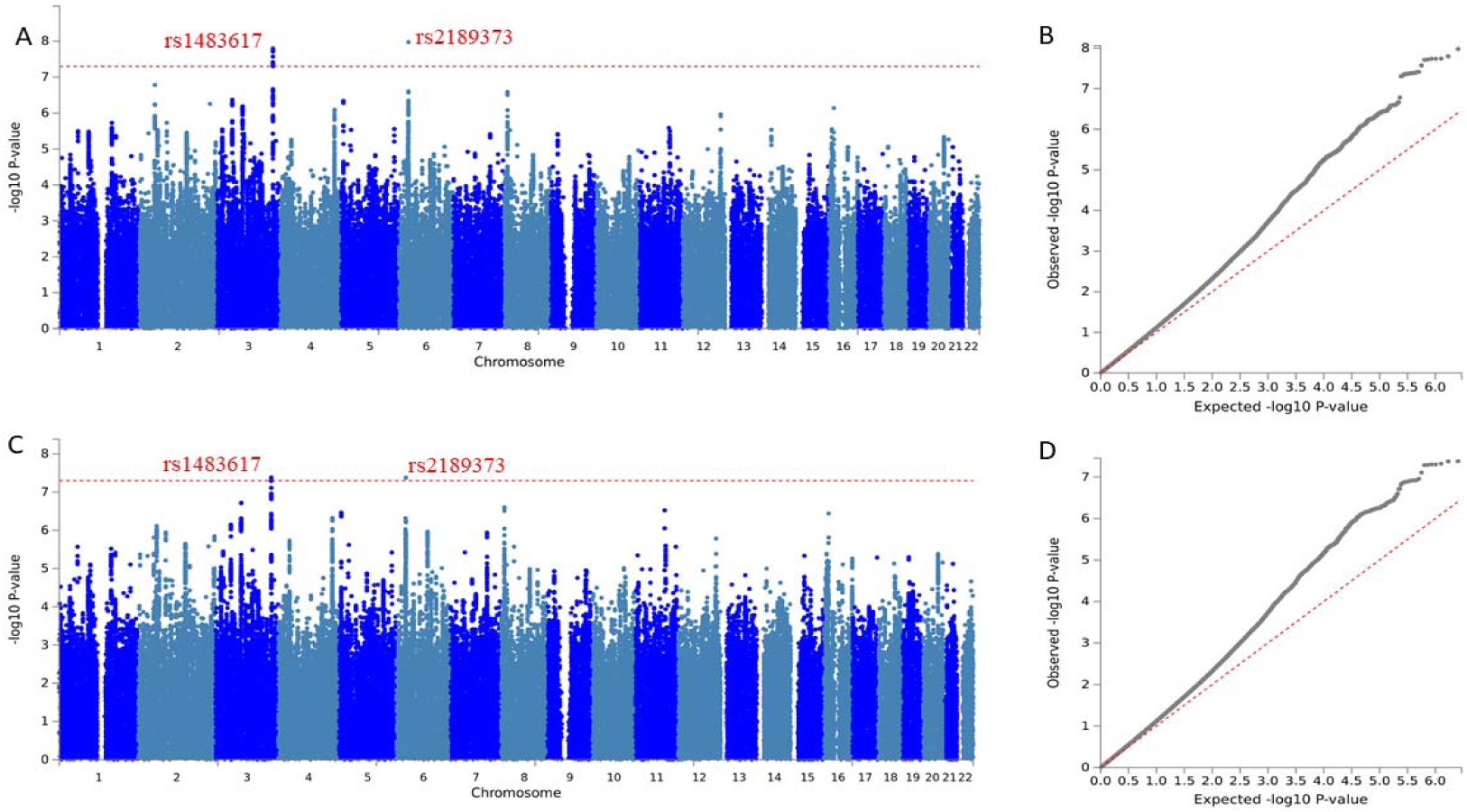
Manhattan and QQ-plots. Manhattan and QQplots of the two MTAG-GWAS datasets. A: Manhattan plot of family relationship satisfaction (N_*effective*_ = 164,112). B: QQplot of family relationship satisfaction (LDSR intercept = 0.99(0.0084); λ_*GC*_*= 1.18). C. Manhattan plot for friendship satisfaction N*_*effective*_ = 158,116). D QQplot for friendship satisfaction (LDSR intercept = 0.99(0.0087); λ_*GC*_= 1.17). Significant SNPs have been labelled on the Manhattan plots in red.

**Figure 3:**
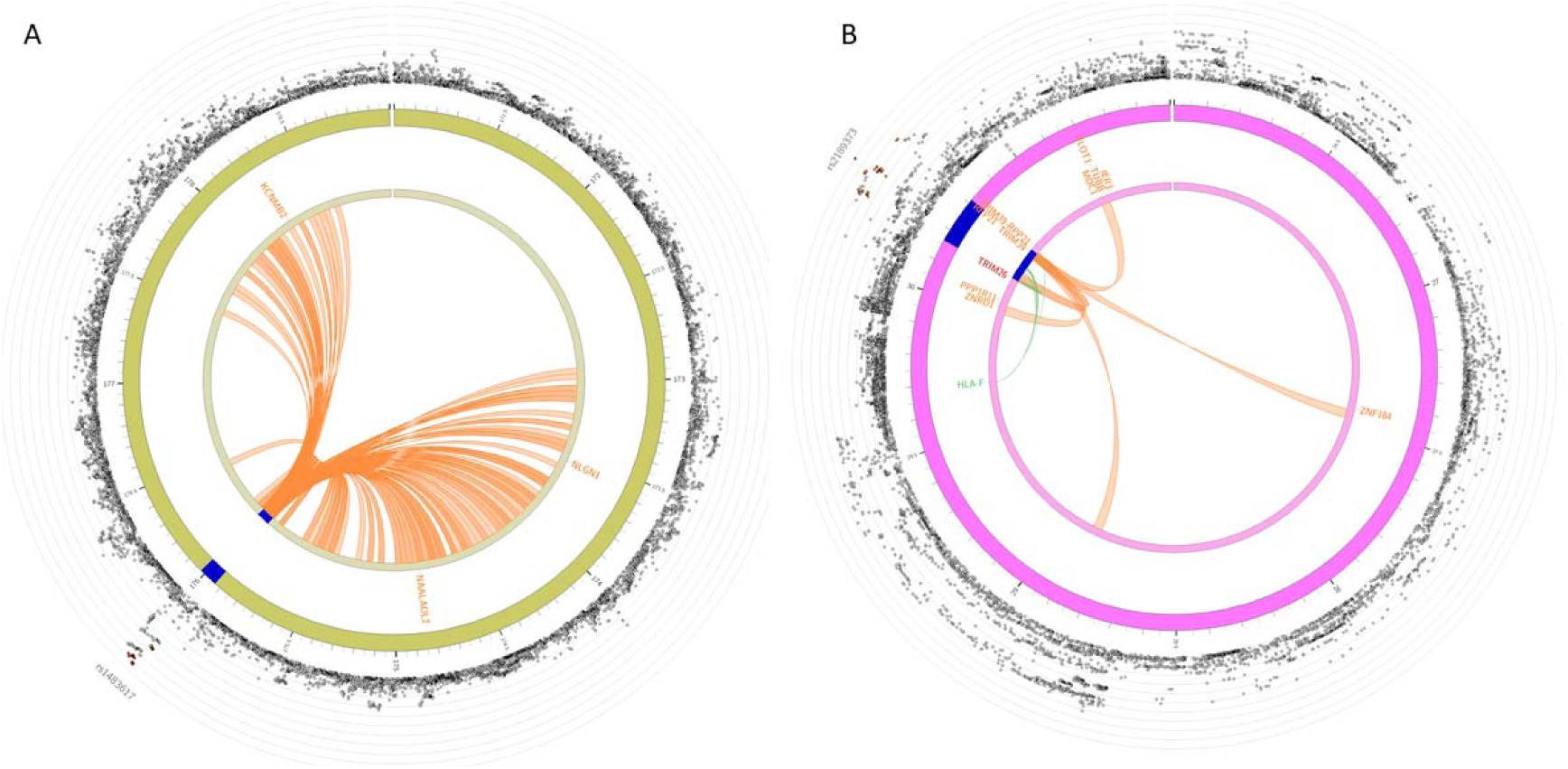
Circos plots showing chromosome interactions and eQTLs. Circos plots showing eQTLs (green lines) and chromosome interactions (orange lines) for the two significant loci. A: Circos plot for rs1483617. B: Circos plot for rs2189373. The outer ring shows a modified Manhattan plot with −log10 P-values. Blue regions identify genomic risk regions. Names of genes with significant chromosome interactions or eQTL interactions are provided in the middle circle. Red regions indicate regions implicated by both eQTL and chromosome interactions.

### Genetic correlation

Despite the modest phenotypic correlation, genetic correlation between family relationship satisfaction and friendship satisfaction was high (r_g_ = 0.87±0.03; P < 2.2x10^-16^), suggesting a similar genetic architecture between the two phenotypes. We observed a similarly high bivariate genetic correlation using GCTA REML (r_g_ = 1 ± 0.11; P < 2.2x10^-16^) in our subsample of participants. This similarity was reflected in the largely similar genetic correlation between the two phenotypes and other psychiatric and psychological phenotypes. After FDR correction, we identified significant negative genetic correlations between family relationship satisfaction and six of the seven psychiatric conditions tested (anxiety, autism, anorexia nervosa, bipolar disorder, depression, and schizophrenia) and between friendship satisfaction and four of the seven psychiatric conditions (autism, bipolar disorder, depression, and schizophrenia). We replicated the negative genetic correlations with autism using a separate autism GWAS dataset (autism-iPsych). Underscoring the contribution to the two phenotypes to most psychiatric conditions, we also identified significant negative genetic correlation for both phenotypes and cross-disorder psychiatric GWAS. For cognitive and psychological phenotypes, we identified negative genetic correlations for both phenotypes and educational attainment, cognitive aptitude, depressive symptoms and neuroticism. In contrast, we identified significant positive genetic correlations between both phenotypes and conscientiousness, empathy, extraversion, and subjective wellbeing. Genetic correlations are provided in **Figure 1 and Supplementary Table 1**.

### SNP heritability

LDSR based SNP-heritability was 0.065 (0.005) for friendship satisfaction and 0.066 (0.005) for family relationship satisfaction. GCTA based heritability for a subset of the participants identified a significant heritability of 0.053 (0.014) for friendship satisfaction and 0.056 (0.014) for family relationship satisfaction (**Supplementary Figure 5 and Supplementary Table 2**).

### Genetic association analyses

Genome-wide association analyses did not identify any significant results. Manhattan and QQ-plots are provided in **Supplementary Figure 6**. The λ_GC_ was 1.15 for both phenotypes. However, the LDSR intercept was 1.0018 (0.007) for family relationship satisfaction and 1.0002 (0.0085) for friendship satisfaction, suggesting negligible inflation due to population stratification. The high genetic correlation between the two phenotypes was supported by concordant effect direction for all the 13 independent loci with P < 1x10^-6^ for the two phenotypes (**Supplementary Figure 7**). Six of these had a significant P-values in the other phenotype after Bonferroni correction (**Supplementary Figure 7 and Supplementary Table 3**).

Given the similar mean chi-square of the two phenotypes (1.13 for family relationship satisfaction and 1.14 for friendship satisfaction) and the high genetic correlation, we conducted a modified inverse variance weighted meta-analysis (MTAG). This increased the effective sample size to 164,112 for family relationship satisfaction and over 158,166 for friendship satisfaction. There was a high genetic correlation between the pre-MTAG and post-MTAG GWAS datasets (Family: 0.98±0.005, P < 2.2x10^-16^; Friendship: r_g_ = 0.98±0.004; P < 2.2x10^-16^). We identified two significant loci associated with both the phenotypes **(Supplementary Table 4)**. On Chromosome 3, the lead SNP was rs1483617 for both phenotypes (P = 1.6x10^-8^ for family relationship satisfaction and 4.55x10^-8^ for friendship satisfaction). In addition, on chromosome 6, we identified another locus with rs2189373 as the lead SNP for both phenotypes (P = 1.06x10^-8^ for family and P = 4.21x10^-8^ for friendship). The top-SNPs explained 0.019% of the variance, which reduced to 0.0027% after correcting for winner’s curse. Manhattan and QQ-plots for the GWAS are provided in **Figure 2.**

### Functional annotation

We investigated genes associated with the two loci using eQTL and chromosome interaction analysis in brain tissues **(Supplementary Tables 5 and 6)**. The analyses prioritized several genes including *NLGN1* which is involved in the postysynaptic complex in excitatory synapses (rs1483617) and interacts with *NRXN1*, a gene implicated in schizophrenia^31,32^, autism^33^, and intellectual disability^34^. A second gene identified, *KCNMB2*, modulates the calcium sensitivity in the BK channels. The second locus on chromosome 6 (rs2189373) is in the MHC complex. It is an intergenic SNP in *HCG17*, a gene which has been implicated in schizophrenia^35^. SNPs in high LD (r^2^ > 0.8) with the lead SNP have been implicated in schizophrenia (rs2021722, P = 2.2 × 10^-12^)^36^ (rs2523722, P = 1.47x10^-16^)^37^ and in the cross-disorder psychiatric GWAS (rs2517614, P = 8.9x10^-7^)^38^. When comparing the two previous studies and our current study, the effect allele which increased the risk for the psychiatric condition is in the same haplotype as the effect allele which decreased social relationship satisfaction. We note here that this region was not genotyped in the latest GWAS of schizophrenia^39^, and no SNP in r^2^ > 0.8 with rs2189373 was available in this schizophrenia summary statistics. eQTL and CCC identified several genes in this locus. Notably, one of the genes identified (*ZNF184*) was also prioritized in a recent large-scale GWAS on neuroticism^40^. **Figure 3** provides the circos plots for the two loci. **Supplementary Figures 8 and 9** provide the local LD plots for the two phenotypes

Using two different methods (FUMA and LDSR-partitioned heritability), we identified high enrichment in brain tissues including the pituitary, pointing to the significant role of neural tissues for both phenotypes **(Supplementary Figures 10 and 11, Supplementary Tables 7 - 9)**. Brain specific chromatin modifications showed a significant, 3-fold enrichment using LDSR, accounting for nearly a third of the heritability **(Supplementary Tables 7 and 8)**. In addition, both methods identified significant enrichment for genes with PLI > 0.9 (Family: Partitioned-h^2^ P = 6.81x10^-7^, MAGMA P = 3.06x10^-4^; Friendship: Partitioned-h^2^ P = 1.56x10^-6^; MAGMA_P = 9.62x10^-5^) **(Supplementary Tables 7 and 8)**. Baseline partitioned heritability also identified enrichment for conserved regions and H3K4me1 modifications for both phenotypes.

We investigated enrichment in specific brain regions using FUMA and partitioned heritability **(Supplementary Figures 10 and 11, Supplementary Table 9)**. FUMA identified a significant enrichment for the cerebellum. In contrast, partitioned heritability identified a significant enrichment for the cortex and the anterior cingulate cortex. This is likely to be due to methodological differences. Cell type specific analysis did not identify a significant enrichment for any specific brain types, using both partitioned heritability and MAGMA **(Supplementary Tables 10 and 11)**.

Gene-based analyses using MAGMA identified thirteen significant genes for family relationship satisfaction and 12 significant genes for friendship satisfaction (**Supplementary Tables 12 and 13)**. The top gene in both the phenotypes was *SHISA5*, a gene that encodes a transmembrane protein on the endoplasmic reticulum.

### Polygenic score analyses

We identified a high correlation between polygenic scores for the two phenotypes in the ALSPAC cohort (r = 0.93; P < 2.2x10^-16^), which reflects the high genetic correlation between the traits **(Supplementary Figure 12)**. Polygenic prediction analyses were conducted for three childhood questionnaires and 5 subscales (Methods). After FDR correction, polygenic scores did not significantly predict variance in any of the phenotypes **(Supplementary Table 14)**. The variance explained was small, with the highest variance explained being for childhood prosocial behaviour **(Supplementary Table 14 and Supplementary Figure 13 and 14)**.

## Discussion

We present the results of a large-scale genome-wide association study of social relationship satisfaction in the UK Biobank measured using family relationship satisfaction and friendship satisfaction. Despite the modest phenotypic correlations, there was a significant and high genetic correlation between the two phenotypes, suggesting a similar genetic architecture between the two phenotypes.

We first investigated if the two phenotypes were genetically correlated with psychiatric conditions. As predicted, most if not all psychiatric conditions had a significant negative correlation for the two phenotypes. We replicated the correlation with autism using a second dataset. We observed significant negative genetic correlation between the two phenotypes and a large cross-condition psychiatric GWAS^38^. This underscores the importance of social relationship dissatisfaction in psychiatric conditions. The genetic correlations identified here are very similar to those identified between psychiatric conditions, psychological phenotypes and subjective well-being^41^. One notable exception is the negative genetic correlation between measures of cognition^42,43^ and the two phenotypes. Whilst subjective wellbeing is positively genetically correlated with measures of cognition, we identify a small but statistically significant negative correlation between measures of correlation and the two phenotypes. A recent study highlighted that people with very high IQ scores tend to report lower satisfaction with life with more frequent socialization^44^. This also highlights the distinctions between subjective wellbeing and social relationship satisfaction. Subjective wellbeing encompasses several aspects of wellbeing including social relationships satisfaction.

We leveraged the high genetic correlation between the two phenotypes to identify significant genes using a method known as MTAG^12^. MTAG makes several assumptions, the key assumption being that the shared variance-covariance matrix is uniform across all SNPs. Despite this, we were comfortable applying MTAG to the two phenotypes due to both the high genetic correlation measured using two different methods, and the similar statistical power of the two GWAS, identified using the similar mean chi-square.

Our MTAG GWAS identified two significant loci. The first (rs1483617) is, as far as we are aware, a novel locus that has not been implicated previously in any GWAS of a neuropsychiatric condition. eQTL and chromosome interactions prioritize several genes that interact with this locus including *NLGN1*, which codes for the post-synaptic protein neuroligin1. NLGNs along with NRXNs and SHANKs are integral component of the synaptic complex, and mutations in these group of proteins have been implicated in several neuropsychiatric conditions including autism^45^. NLGN1 is located in the post-synaptic membrane of glutamatergic synapses, and can modulate the development of glutamatergic synapses in an activity dependent manner^45^. Another gene prioritized at this locus, *KCNMB2*, encodes a protein that is a component of the BK channels which are large conductance potassium channels that are sensitive to both intracellular voltage changes and changes in calcium ions.

The second locus (rs2189373) is in the MHC, a region of complex LD structure. This locus has been implicated previously in schizophrenia, and is nominally associated with the cross-condition psychiatric GWAS. Interestingly, this locus was not investigated in the recent large-scale GWAS of schizophrenia^39^, as inferred from the summary statistics available. The effect direction of the SNPs when considering the haplotype structure of this region align with the genetic correlation between social relationship satisfaction and psychiatric risk. In other words, alleles that increase the risk for schizophrenia are in the same haplotype as alleles that decrease friendship satisfaction. The functional consequences of this locus must be formally tested.

Multiple different lines of evidence also point to the central role of brain tissues for the two phenotypes. First, using partitioned heritability, we identified a significant, threefold-enrichment in brain specific annotations for both phenotypes, accounting for about 30% of the total heritability. This mirrored the high enrichment in brain specific expression identified by FUMA, including the pituitary. We also sought to identify where within the brain tissue GWAS signals for the two phenotypes are enriched in. Here, FUMA and partitioned heritability identified enrichment in different tissues. While FUMA prioritized the cerebellum, heritability prioritized the cortex. The discordance in results is due to the different definition of tissue specific expression used by the two methods. While partitioned heritability compares focal brain region expression against other brain regions, FUMA compares the focal brain region’s expression against all tissue types. Hence, partitioned heritability is more likely to tag brain region specific expression patterns.

In addition to enrichment for brain tissues, we also identified an enrichment for genes that are intolerant to loss of function mutations^23^. A fifth of the heritability can be attributed to pLI > 0.9 genes. We utilized pLI quantified from the non-psychiatric population cohort for this analysis given the considerable genetic correlation between the two phenotypes and various psychiatric conditions. Loss of function mutations in these genes lead to severe biochemical consequences, and are implicated in several neuropsychiatric conditions. For example, *de novo* loss of function mutations in pLI intolerant genes confers significant risk for autism^46^. Our results suggest that pLI > 0.9 genes contribute to psychiatric risk through both common and rare genetic variation.

Whilst we were unable to investigate the polygenic prediction in other samples with same phenotypic measures, we investigated polygenic prediction across a range of different phenotypes in children. We chose three specific questionnaires that have been associated with various psychiatric conditions^47–49^, and subdomains in the SDQ. After FDR correction, we did not identify any significant polygenic prediction for the two phenotypes. The variance explained was small. This is due to three reasons – the modest genetic correlation between social relationship satisfaction and the childhood phenotypes, the modest statistical power of this study due to both low per-SNP variance explained and low additive heritability, and the relatively small sample size of the ALSPAC cohort. Encouragingly, the phenotype for which the variance explained was the highest is childhood prosocial behaviour.

In conclusion, we identify two significant loci association with social relationship satisfaction, one of which has been previously implicated in schizophrenia. Comprehensive functional annotation prioritizes specific genes for functional follow-up, and identifies tissue specific enrichment pattern. Genetic correlation analyses highlight the importance of social relationship dissatisfaction in psychiatric conditions, and enrichment in loss-of-function intolerance genes suggest a degree of overlap between rare and common variants in a set of genes for psychiatric conditions. Social relationship dissatisfaction is not just a consequence of having a psychiatric condition because both phenotypes have genetic correlates, meaning they are both valid phenotypes. Future research should test the function of the two significant SNPs (rs1483617 and rs2189373) we identified, and their role in each psychiatric condition.

## Acknowledgements

We thank Patrick Turley and Beate St. Pourcain for their helpful discussion on the methods. We thank Anders Børglum, Jakob Grove, and the iPSYCH team for generously sharing the data with us. We thank 23andMe for giving us access to summary statistics for this study, and the research volunteers at 23andMe for making this research possible. The 23andMe Research Team are: Michelle Agee, Babak Alipanahi, Adam Auton, Robert K. Bell, Katarzyna Bryc, Sarah L. Elson, Pierre Fontanillas, Nicholas A. Furlotte, David A. Hinds, Karen E. Huber, Aaron Kleinman, Nadia K. Litterman, Jennifer C. McCreight, Matthew H. McIntyre, Joanna L. Mountain, Elizabeth S. Noblin, Carrie A.M. Northover, Steven J. Pitts, J. Fah Sathirapongsasuti, Olga V. Sazonova, Janie F. Shelton, Suyash Shringarpure, Chao Tian, Joyce Y. Tung, Vladimir Vacic, and Catherine H. Wilson. We are grateful to all the families who took part in ALSPAC, the midwives for their help in recruiting them, and the whole ALSPAC team, which includes interviewers, computer and laboratory technicians, clerical workers, research scientists, volunteers, managers, receptionists and nurses. VW is funded by St. John’s College, Cambridge, and the Cambridge Commonwealth Trust. This study was funded by grants from the Medical Research Council, the Wellcome Trust, the Autism Research Trust, the Templeton World Charity Foundation, the Institut Pasteur, the CNRS and the University Paris Diderot. The research was funded and supported by the National Institute for Health Research (NIHR) Collaboration for Leadership in Applied Health Research and Care East of England at Cambridgeshire and Peterborough NHS Foundation Trust. We acknowledge with gratitude the generous support of Drs Dennis and Mireille Gillings in strengthening the collaboration between SBC and TB, and between Cambridge University and the Institut Pasteur. The views expressed are those of the author(s) and not necessarily those of the NHS, the NIHR or the Department of Health. Data obtained from 23andMe was supported by the National Human Genome Research Institute of the National Institutes of Health (grant number R44HG006981). The UK Medical Research Council and Wellcome (grant ref: 102215/2/13/2) and the University of Bristol provide core support for ALSPAC. GWAS data was generated by Sample Logistics and Genotyping Facilities at Wellcome Sanger Institute and LabCorp (Laboratory Corporation of America) using support from 23andMe. This publication is the work of the authors who will serve as guarantors for the contents of this paper.

## Supplementary Note

### Index

#### Supplementary Figures

S1: Phenotypic distributions of family and friendship relationship satisfaction (*3*)

S2: Spearman’s rank correlation between phenotypic distributions of friendship and family relationship satisfaction (*4*)

S3: Difference in scores for friendship and family relationship satisfaction based on sex (*5*)

S4: Difference in scores for friendship and family relationship satisfaction based on age (*6*)

S5: Additive SNP heritability for family relationship and friendship satisfaction (*7*)

S6: Manhattan and QQ-plot for pre-MTAG GWAS (*8*)

S7: Direction and P-values of all independent SNPs with P < 10^-6^ in the pre-MTAG family relationship and friendship satisfaction GWAS (*9*)

S8: Regional LD plot for family relationship satisfaction (*10*)

S9: Regional LD plot for friendship satisfaction (*11*)

S10: General tissue enrichment (FUMA) (*12*)

S11: Specific tissue enrichment (FUMA) (*13*)

S12: Correlation in Polygenic scores (*14*)

S13: Distribution of phenotypes tested in polygenic regression analysis (*15)*

S14: Polygenic regression analyses (*16*)

Supplementary Note 1: ALSPAC cohort profile (*17*)

Supplementary Note 2: Proxy-phenotype replication (*18*)

**Supplementary Figure 1:**
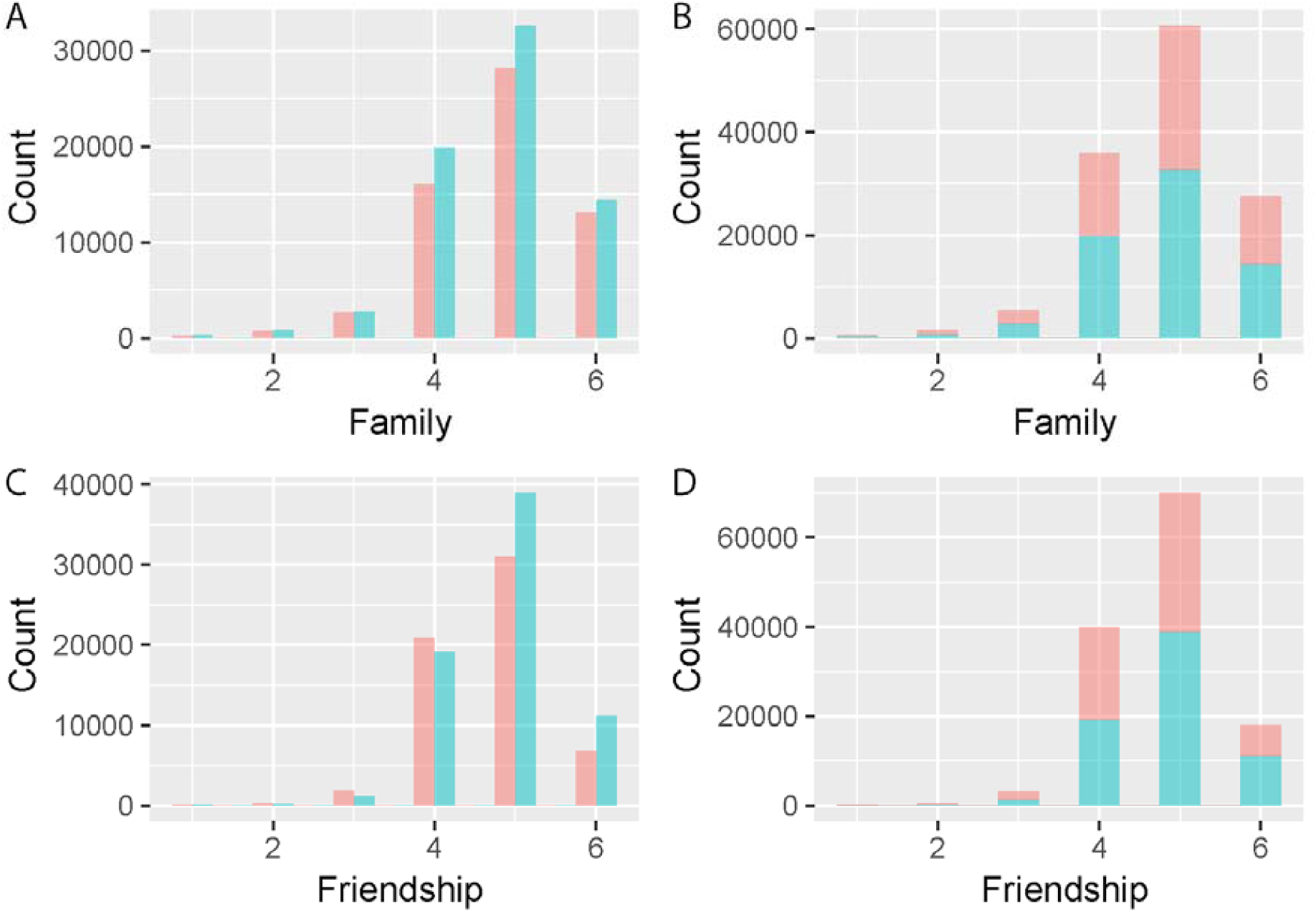
Phenotypic distributions of family and friendship relationship satisfaction. Histograms plotting the distribution of family relationship satisfaction (A and B), and friendship satisfaction (C and D). Blue bar indicates scores for females, and pink for males. Plots B and D show stacked frequency histograms. Scores range from 1 to 6, with 1 corresponding to extremely unhappy and 6 corresponding to extremely happy.

**Supplementary Figure 2:**
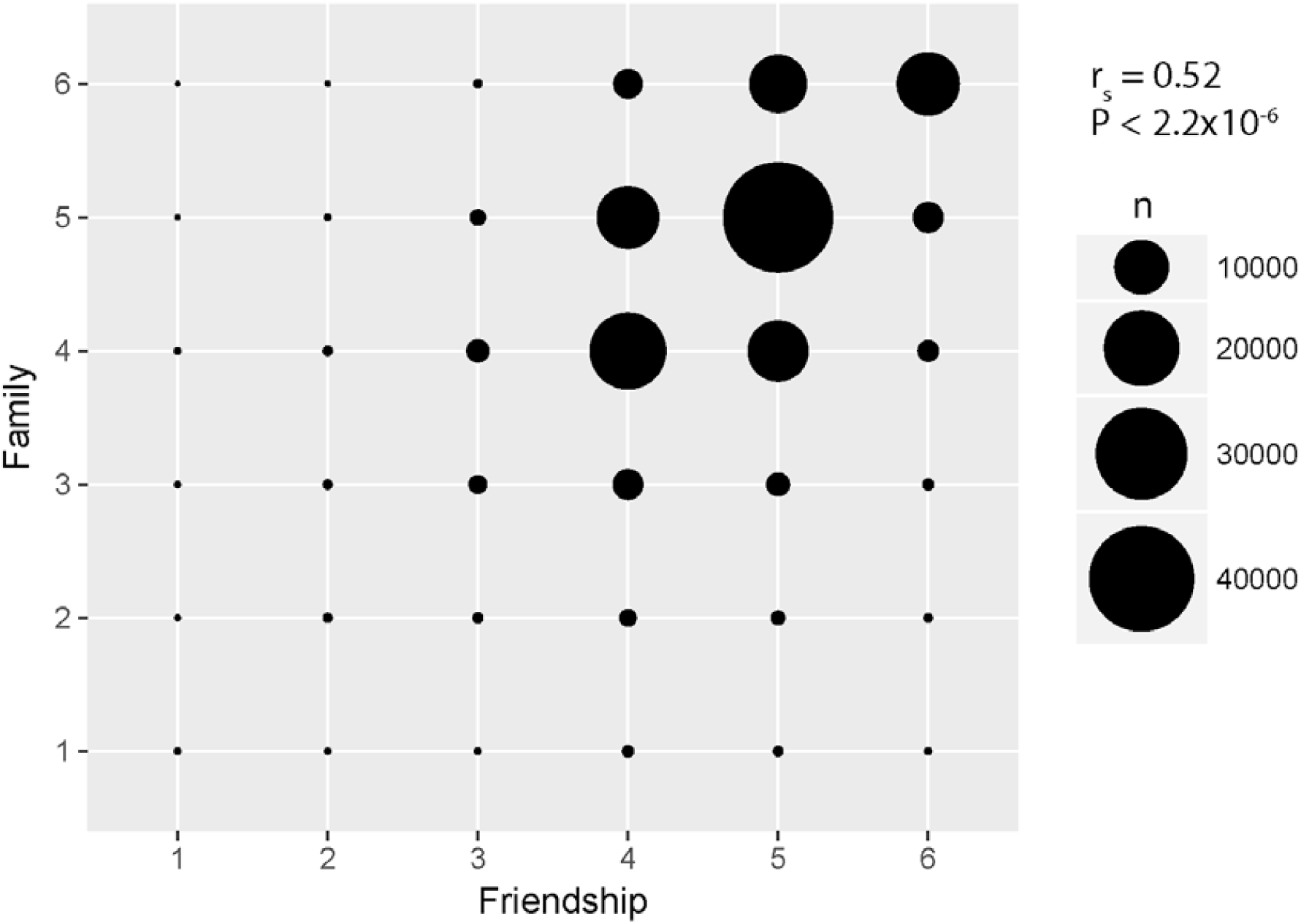
Spearman’s rank correlation between phenotypic distributions of friendship and family relationship satisfaction. Each circle shows the overlap between scores on the two phenotypes. Larger the circle, larger the overlap. Spearman’s rank correlation = 0.52 (P < 2.2x10-6).

**Supplementary Figure 3:**
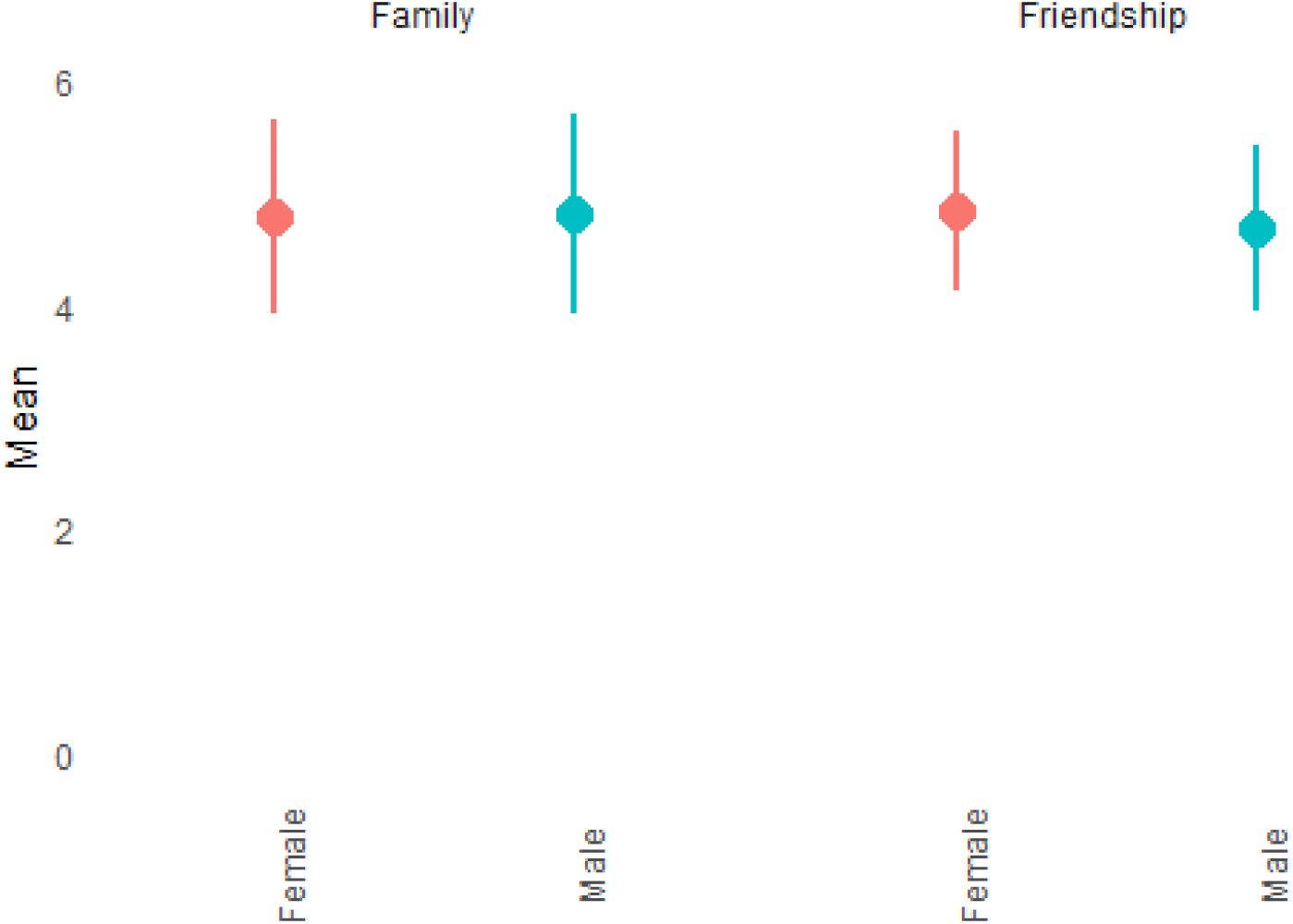
Difference in scores for friendship and family relationship satisfaction based on sex. Mean scores for family relationship (left) and friendship (right) satisfaction shown above. Lines represent standard deviations. There was a small, but significant difference between males and females (Unpaired T-test; P < 0.001 for both the phenotypes). For family relationship satisfaction, males scored significantly higher than females. Mean (males) = 4.81, sd = 0.89; Mean (females) = 4.79; sd = 0.87. The Cohen’s D for the T-test was small (0.02). For friendship satisfaction, males scored significantly lower than females. Mean (males) = 4.68, sd = 0.74; Mean (females) = 4.84, sd = 0.71. The Cohen’s D for the T-test was larger than that of family relationship satisfaction, but still small (0.22). The test was conducted in N = 131790 individuals who completed both the questionnaires. N = 70809 for females. N = 60981 for males.

**Supplementary Figure 4:**
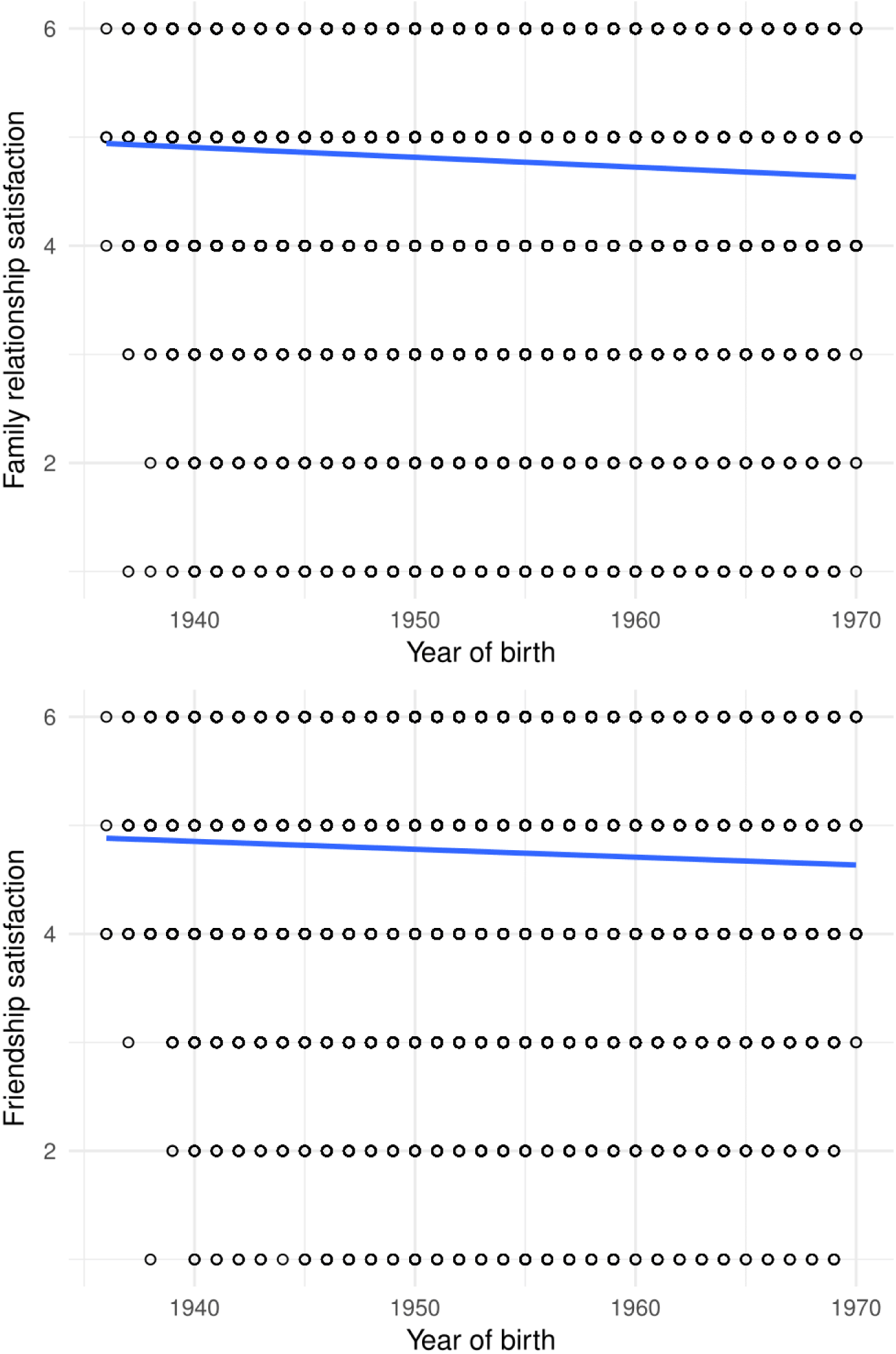
Difference in scores for friendship and family relationship satisfaction based on age. Scatterplot showing the relationship between year of birth and family relationship satisfaction (above) and friendship satisfaction (below). Age predicted a small proportion of the variance in family satisfaction (Beta = −0.009; s.e. = 0.0003 P < 2.2x10-16). Age also predicted a small proportion of the variance in friendship (Beta = −0.007; s.e. = 0.0002; P < 2.2x10-16).

**Supplementary Figure 5:**
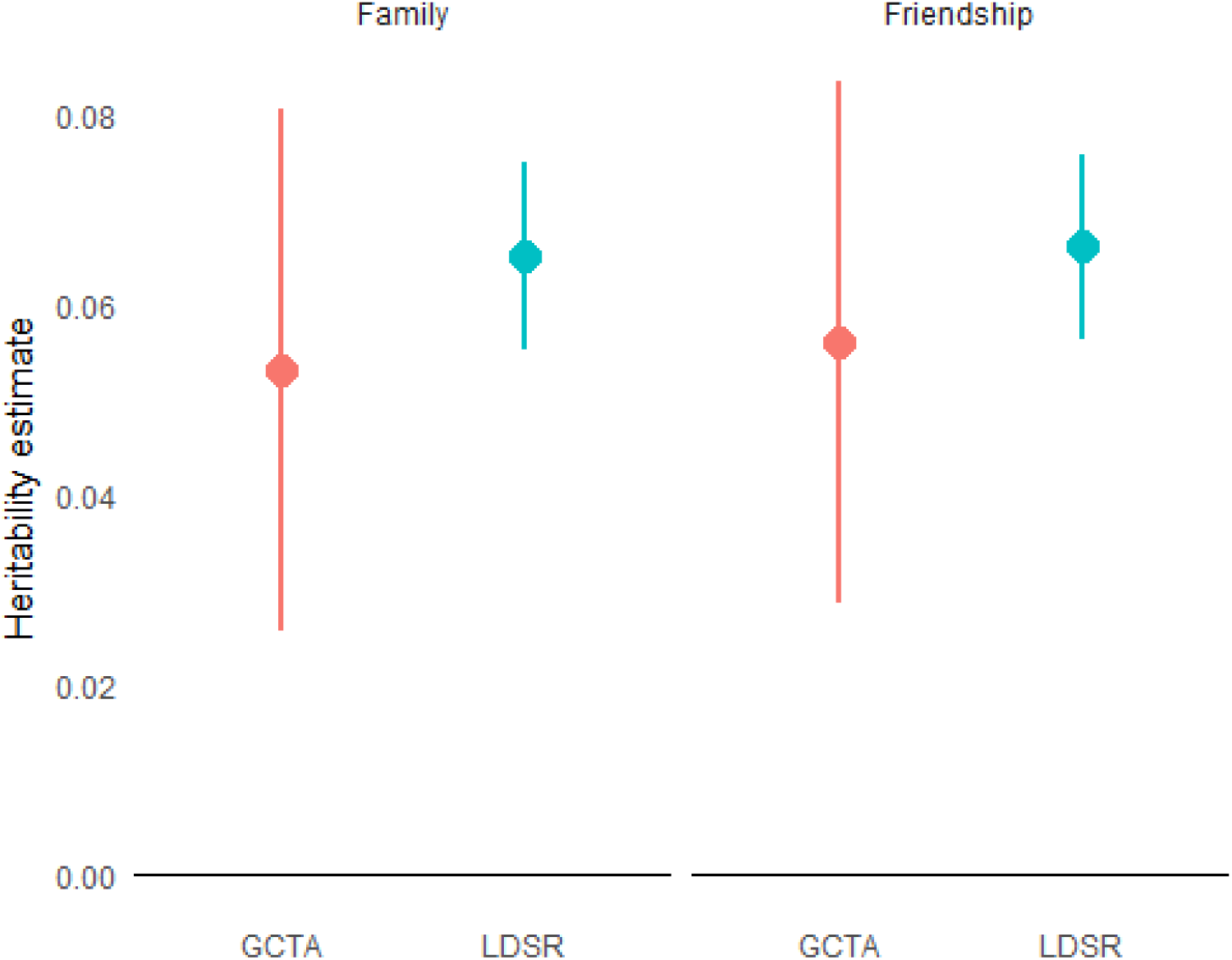
Additive SNP heritability for family relationship and friendship satisfaction. Additive SNP heritability for family relationship satisfaction (left) and friendship satisfaction (right). Heritability estimates and 95% confidence intervals provided. We used two different methods to calculate additive SNP heritability: LD score regression and GCTA genomic-relatedness-based restricted maximum-likelihood. Heritability estimates for the former was calculated using the full sample (N > 130,000 for both phenotypes). Heritability estimates for the latter was calculated using one-fifth of the full sample (N = 26,000 for both phenotypes), for computational efficiency. For both methods we included sex, batch, year of birth and the first forty genetic principal components as covariates. Heritability estimates were similar and significant for both methods. The heritability estimates are: Family relationship: LDSR - 0.065±0.005, P < 0.001; GCTA - 0.053±0.014, P< 0.001. Friendship: LDSR - 0.066±0.005, P < 0.001; GCTA - 0.056±0.014, P< 0.001.

**Supplementary Figure 6:**
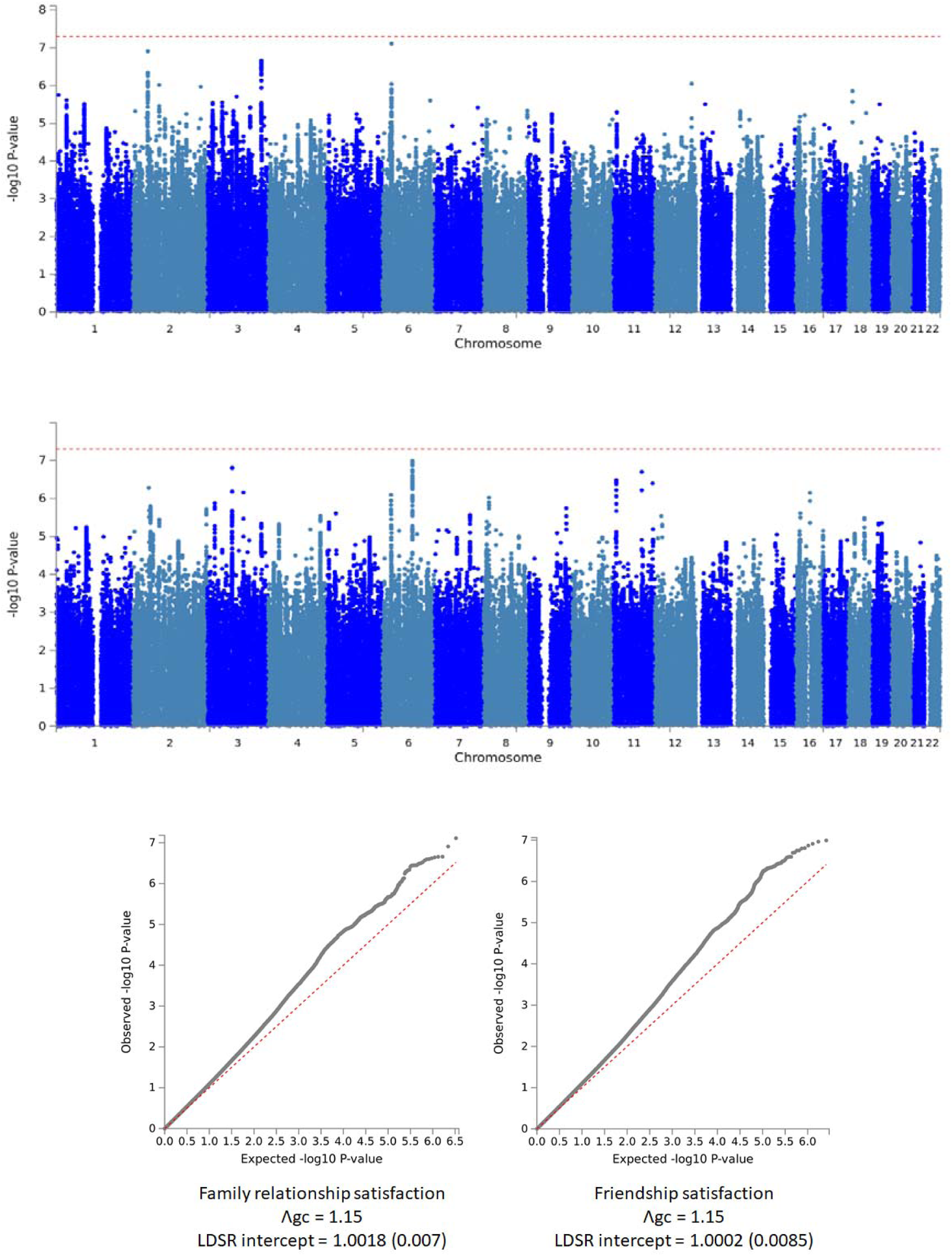
Manhattan and QQ-plot for pre-MTAG GWAS. Manhattan plots for family relationship satisfaction (top), and friendship satisfaction (middle). QQplots for family relationship satisfaction and friendship satisfaction below.

**Supplementary Figure 7:**
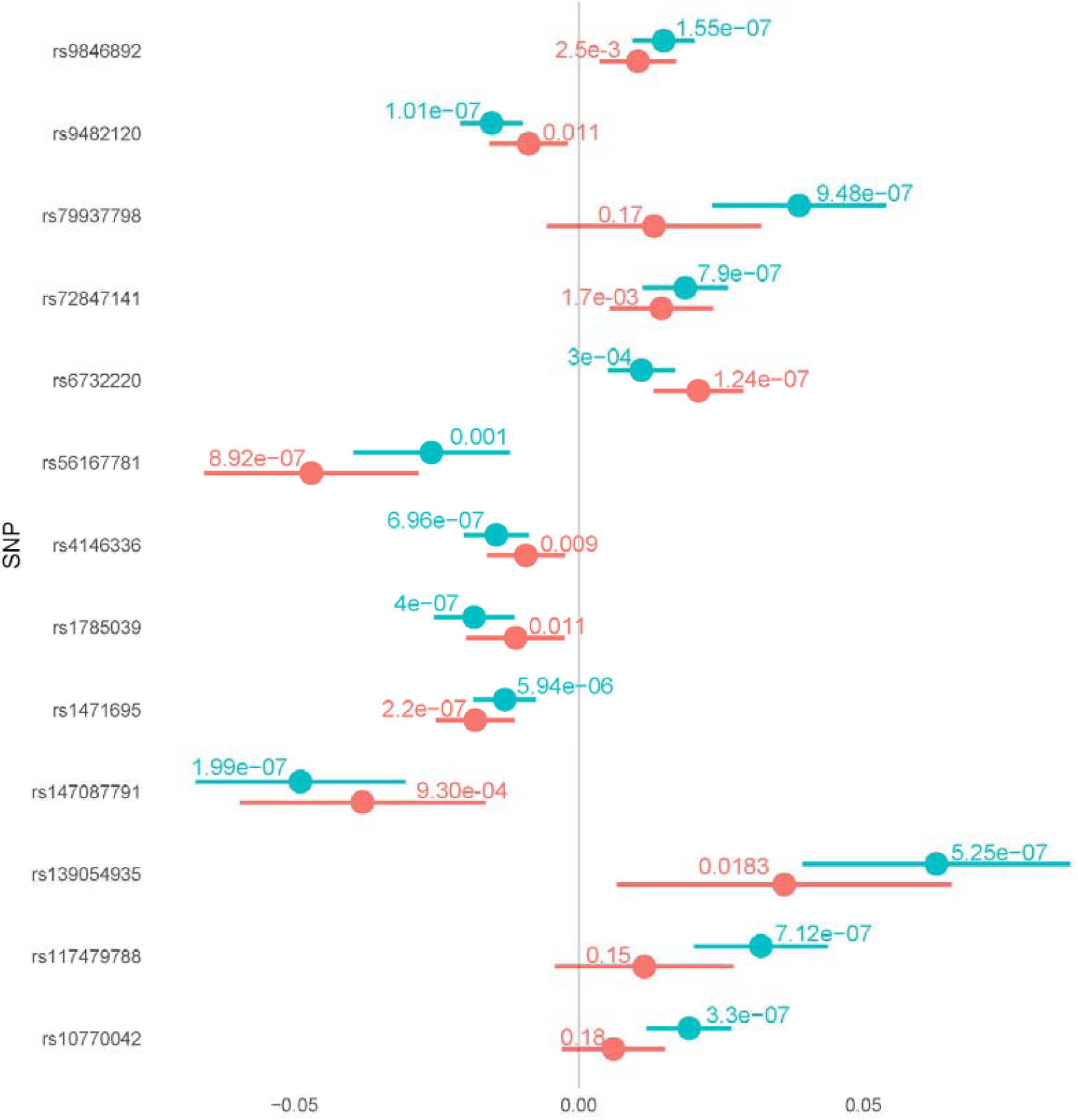
Direction and P-values of all independent SNPs with P < 10^-6^ in the pre-MTAG family relationship satisfaction and friendship satisfaction GWAS. Regression Beta and 95% CI provided for all independent SNPs with P < 1x10-6. Blue lines indicate effect estimate and CI for friendship satisfaction. Red lines indicate effect estimate and 95%CI for family relationship satisfaction. P-values provided on top of the CI lines. We conducted 13 independent tests (replicating 13 independent SNPs). A Bonferroni corrected significant threshold is 0.0038.

**Supplementary Figure 8:**
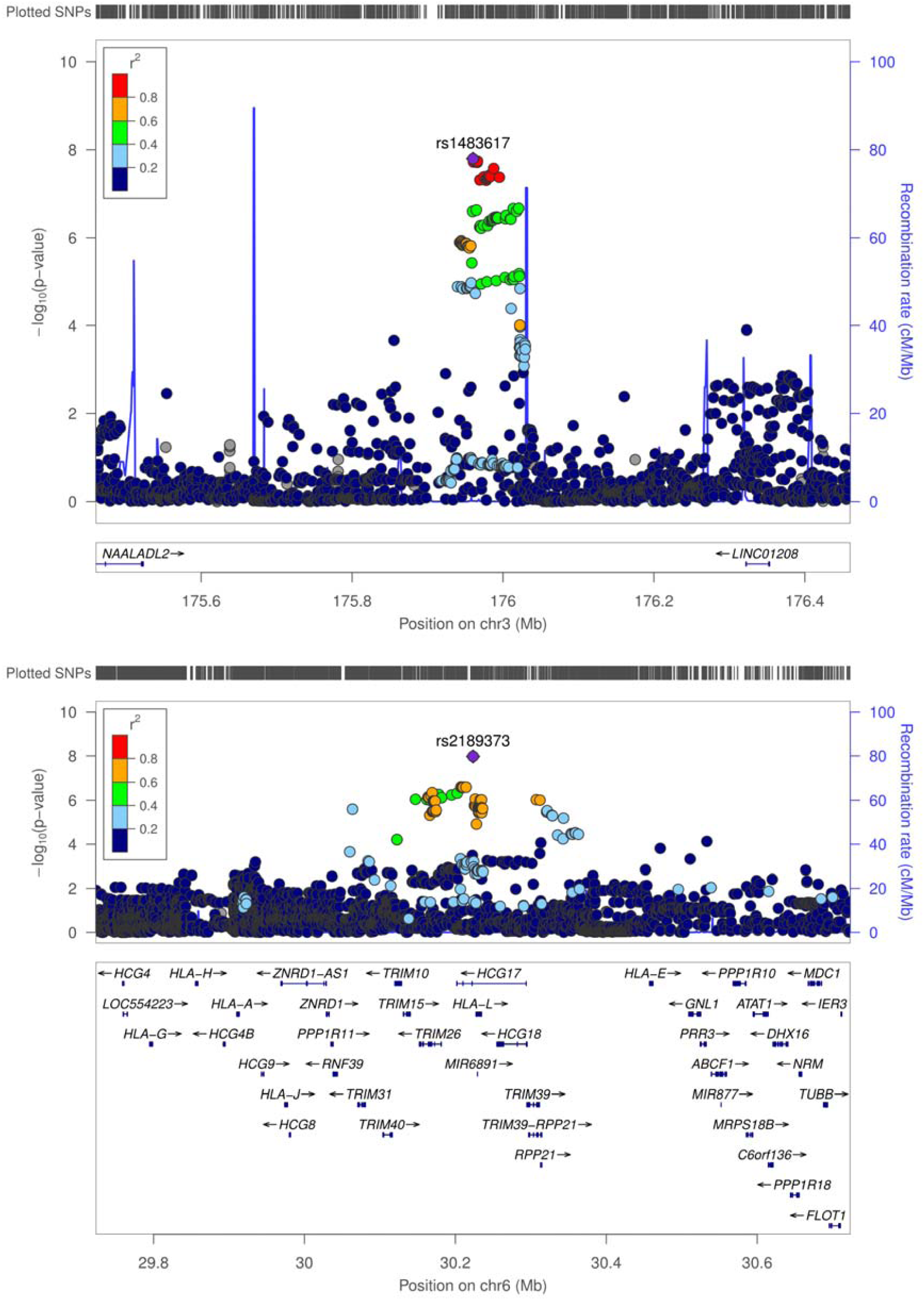
Regional LD plot for family relationship satisfaction.

**Supplementary Figure 9:**
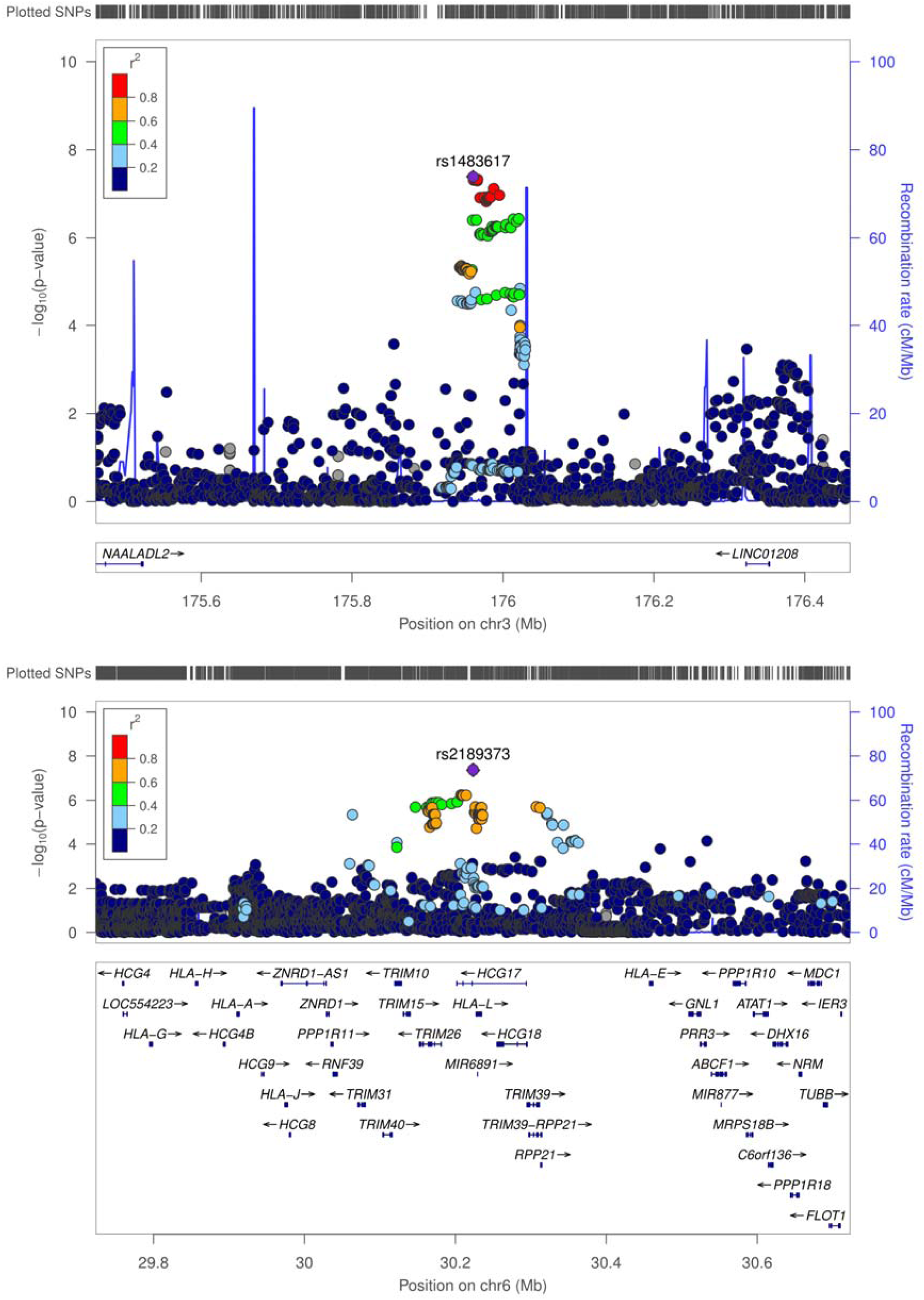
Regional LD plot for the friendship satisfaction.

**Supplementary Figure 10:**
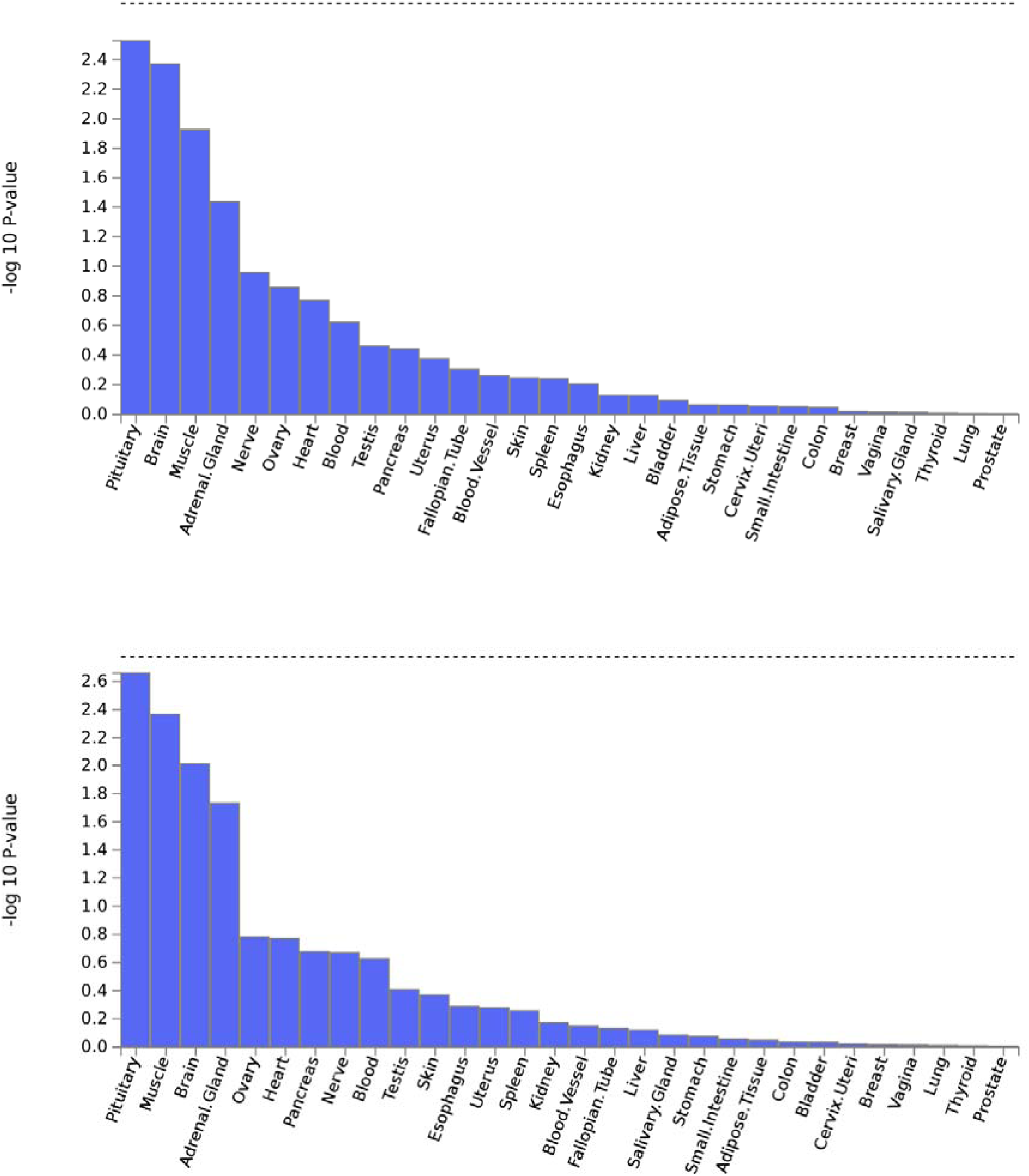
General tissue enrichment (FUMA) FUMA general tissue enrichment plots for family relationship satisfaction (above) and friendship relationship satisfaction (below). X axis indicates tissues, and Y axis indicated enrichment, with higher bars representing greater enrichment. Dotted line indicates Bonferroni corrected threshold. Gene expression data was obtained from the GTEx consortium.

**Supplementary Figure 11:**
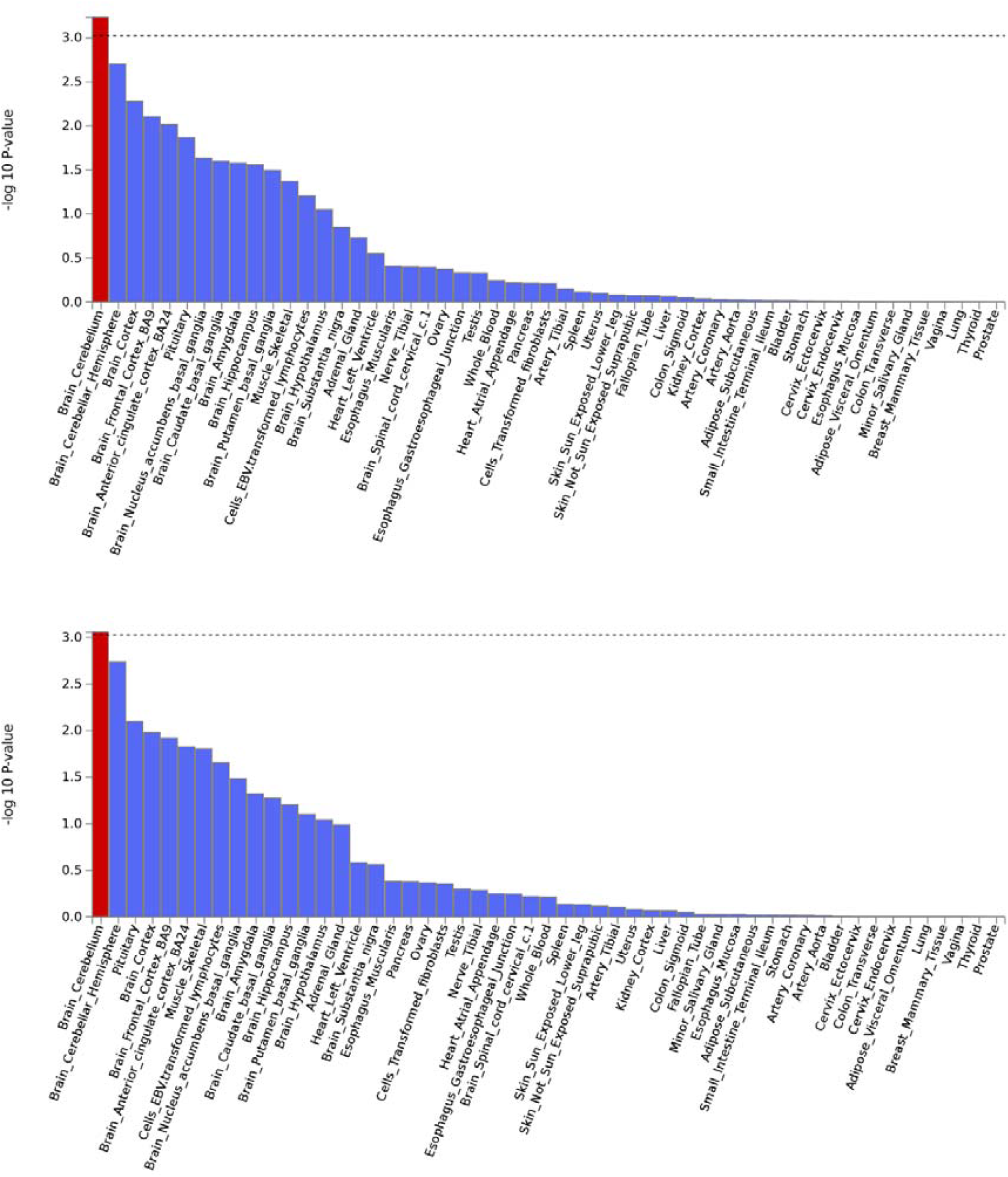
Specific tissue enrichment (FUMA) FUMA specific tissue enrichment plots for family relationship satisfaction (above) and friendship relationship satisfaction (below). X axis indicates tissues, and Y axis indicated enrichment, with higher bars representing greater enrichment. Red bars indicate significant enrichment after Bonferroni correction. Dotted line indicates Bonferroni corrected threshold. Gene expression data was obtained from the GTEx consortium.

**Supplementary Figure 12:**
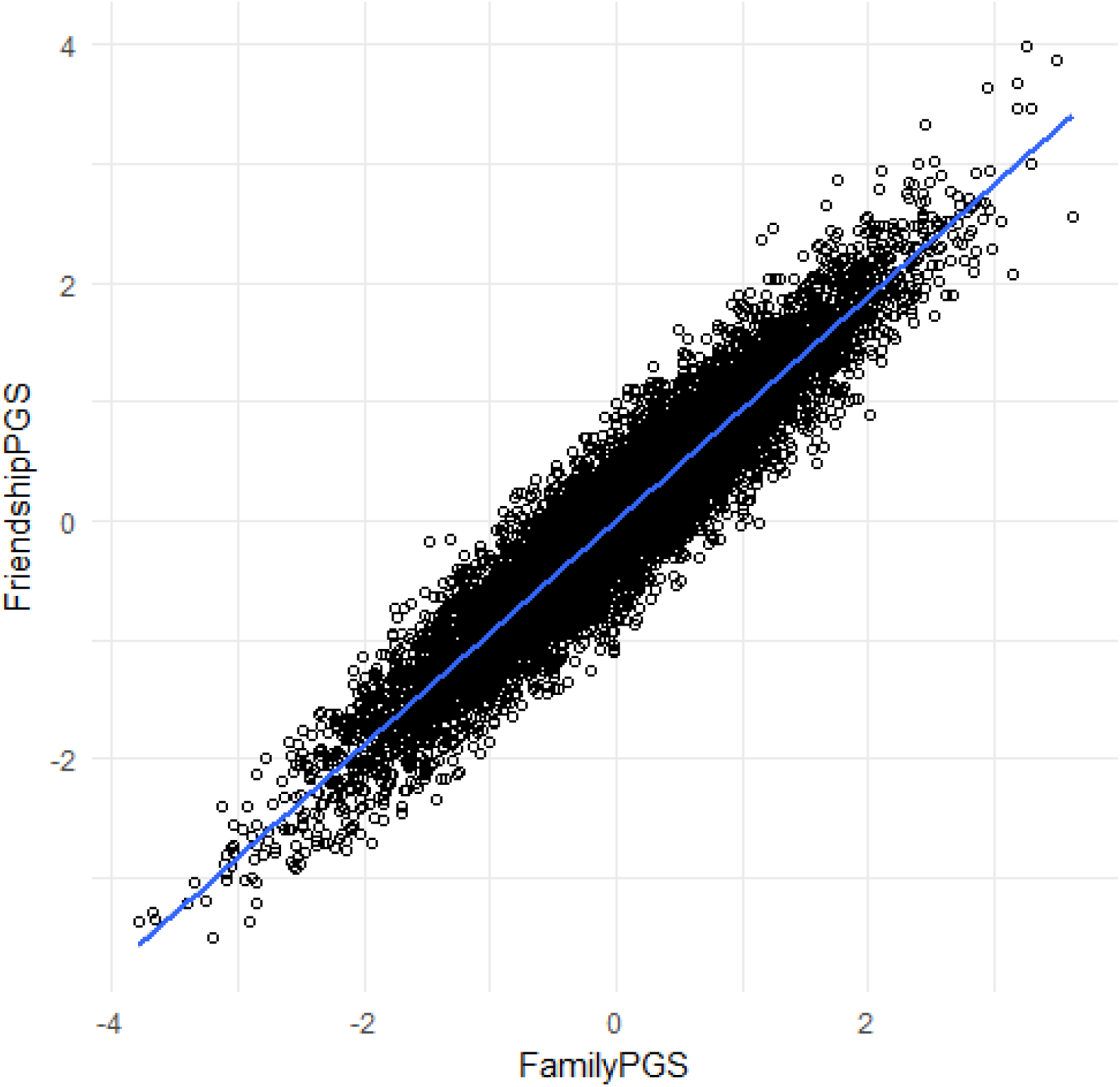
Correlation in Polygenic scores. Pearson’s correlation of scaled polygenic risk scores in 8104 unrelated individuals from ALSPAC. Correlation coefficient = 0.937, 95%CI: 0.934 - 0.939. P < 2.2x10-16

**Supplementary Figure 13:**
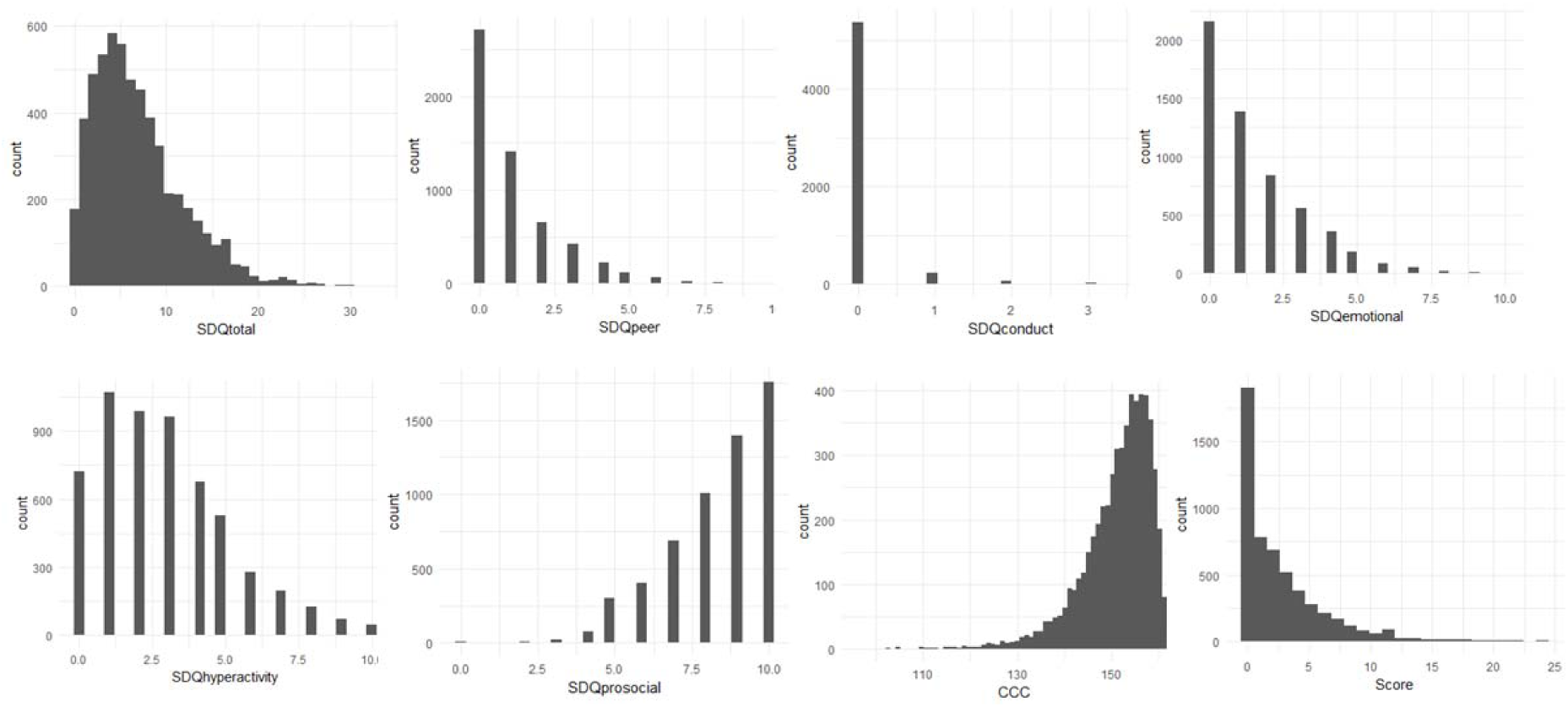
Distribution of phenotypes tested in polygenic regression analysis. Plotting the distribution of all the eight phenotypes revealed non-gaussian distribution, which is expected from count data. All eight phenotypes had poisson distributions with overdispersion. Given the distribution of the traits, polygenic regression analyses were conducted using a negative binomial regression model with sex, and the first two genetic principal components included as covariates. We did not include age as a covariate as the phenotypes were measured at specific ages. In the top row, from left, the phenotypes are: 1. Total SDQ total score (age = 115 months, n = 5646); 2. SDQ peer relationship score (age = 115 months, n = 5660); 3. SDQ conduct problems score (age = 115 months, n = 5650); 4. SDQ emotional symptoms score (age = 115 months, n = 5650). In the bottom row, from left, the phenotypes are: 5 SDQhyperactivity score (age = 115 months, n = 5670); SDQ prosocial score* (age =115 months, n = 5669); 7. Children’s Communication Checklist Score* (age = 115 months, n = 5583); 8. Social and Communication Difficulties questionnaire score (age = 91 months, n = 5447). *These phenotypes were reverse coded in the regression analyses.

**Supplementary Figure 14:**
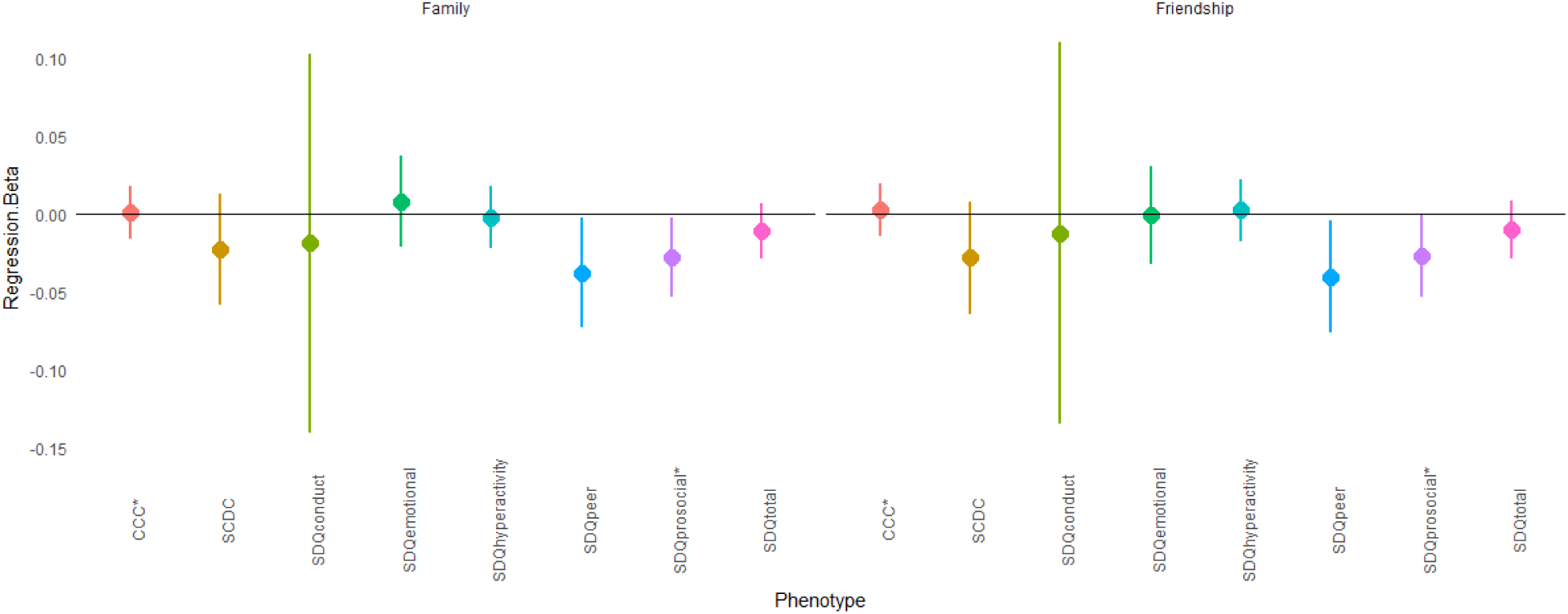
Polygenic regression analyses. Top: Regression beta (effect) of polygenic scores after accounting for sex and the first two genetic principal components for the eight phenotypes tested. Lines indicate 95% confidence intervals. Bottom: Incremental variance explained (Nagelkerke’s pseudo-R2) by the polygenic scores for the eight phenotypes. P-values provided on top.

## Supplementary Note 1: Details about ALSPAC

### Participants

All participants included in the polygenic score analyses were from an ongoing study - the Avon Longitudinal Study of Parents and Children (ALSPAC). We queried the data using a fully searchable data dictionary which is available here: http://www.bristol.ac.uk/alspac/researchers/access/. The ALSPAC cohort comprises 14,541 initial pregnancies from women resident in Avon, UK resulting in a total of 13,988 children who were alive at 1 year of age. Pregnant women were recruited from 1^st^ April 1991 to 31^st^ December 1992. In addition to this, children were enrolled in other phases which is described elsewhere^1^. In total, 713 additional children were enrolled in this study. The study received ethical approval from the ALSPAC Ethics and Law Committee, and written informed consent was obtained from parent or a responsible legal guardian for the child to participate. Assent was obtained from the child participants where possible. In addition to ethical approval from ALSPAC, we obtain ethical approval from the Human Biology Research Ethics Committee at the University of Cambridge.

### Gentoyping

Participants were genotyped using the Illumina HumanHap550 quad chip by 23andMe. GWAS data was generated by Sample Logistics and Genotyping Facilities at Wellcome Sanger Institute and LabCorp (Laboratory Corportation of America) using support from 23andMe. Individuals were excluded based on gender mismatches, high missingness (> 3%), and disproportionate heterozygosity, if they were of non-European descent (CEU) which was assessed using multidimensional scaling and compared with Hapmap II (release 22), if the cryptic relatedness assessed using IBD was greater than 0.1 SNPs were excluded if the per-SNP missingness was more than 5%, if they violated Hardy-Weinberg equilibrium (P < 1x10^-6^), if the MAF < 0.01. In total, there were 526,688 genotped SNPs. Haplotypes were estimated using data from moterhs using SHAPEIT (v2.r644)^2^. Imputation was performed using Impute2 V2.2.2^3^ using the 1000 genomes reference panel (Phase 1, Version 3). Imputed SNPs were excluded from all further analyses if they had a minor allele frequency < 1% and info < 0.8.

### Phenotypes

For all the phenotypes included in the cross-phenotype polygenic risk score analyses, we used the prorated scores as provided in the ALSPAC datasheet. We considered three main phenotypes (CCC, SDQ and SCDC), as they have been associated with psychiatric conditions. For example, both the CCC^4^ and the SCDC^5^ have been linked to autism. Further, the SDQ has been linked to ADHD and conduct problems^6^. In addition to the total SDQ scores, we also considered the SDQ subscales as they capture difficulties in specific domains. Scores on these measures have been provided for multiple different age groups in the ALSPAC. We considered the age group with the highest number of participants for whom phenotypic data was available, which was, for all phenotypes, the youngest age group.

